# “Coarse-grained simulation reveals key features of HIV-1 capsid self-assembly”

**DOI:** 10.1101/040741

**Authors:** John M. A. Grime, James F. Dama, Barbie K. Ganser-Pornillos, Cora L. Woodward, Grant J. Jensen, Mark J. Yeager, Gregory A. Voth

**Author notes:** Corresponding author: Prof. Gregory A. Voth, 5735 S Ellis Ave., Department of Chemistry, The University of Chicago, Chicago, IL 60637.

## Abstract

The maturation of HIV-1 viral particles is essential for viral infectivity. During maturation, many copies of the capsid protein (CA) self-assemble into a capsid shell to enclose the viral RNA. The mechanistic details of the initiation and early stages of capsid assembly remain to be delineated. We present coarse-grained simulations of capsid assembly under various conditions, considering not only capsid lattice self-assembly but also the potential disassembly of capsid upon delivery to the cytoplasm of a target cell. The effects of CA concentration, molecular crowding, and the conformational variability of CA are described, with results indicating that capsid nucleation and growth is a multi-stage process requiring well-defined metastable intermediates. Generation of the mature capsid lattice is sensitive to local conditions, with relatively subtle changes in CA concentration and molecular crowding influencing self-assembly and the ensemble of structural morphologies.

---

Significant morphological changes are required to convert an “immature” virus particle (virion) of HIV-1 into the mature and infectious form ^1-3^. During maturation, the enzymatic cleavage of Gag polypeptide ^4^ releases capsid protein (CA) to self-assemble into a conical lattice structure enclosing the viral RNA (the capsid, Fig. 1). Failure to generate a suitable capsid precludes infectivity ^5-9^ and so the details of capsid generation are of significant interest as a therapeutic target.

**Figure 1.**
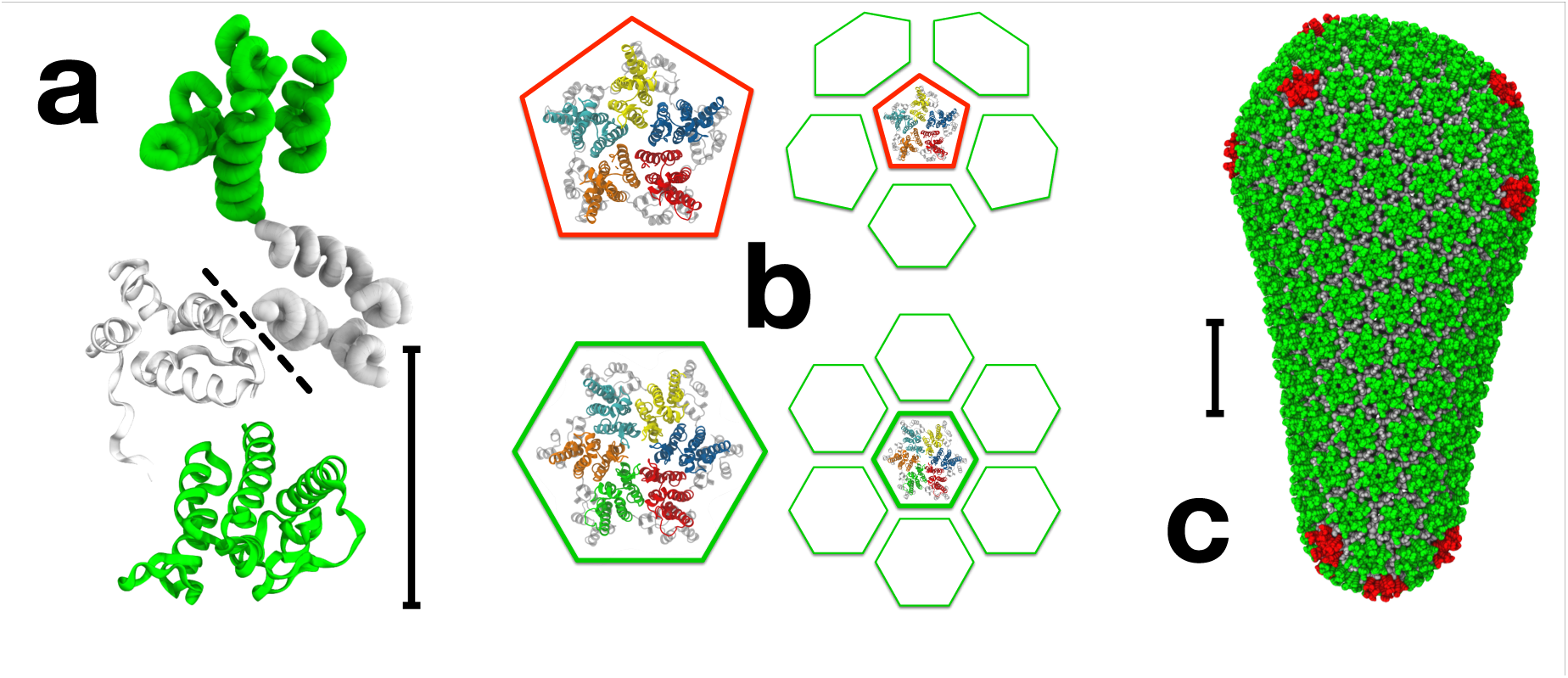
The fullerene cone model of the HIV-1 capsid is assembled from “mature-style” capsid lattice with hexameric (green) and pentameric (red) building blocks. (**a**) CA dimer structure with both CG (tube) and all-atom (ribbon) monomer representations (NTDs green, CTDs gray). CTD dimer interface marked by dashed line, scale bar 5 nm. (**b**) Quasi-equivalent pentamer and hexamer capsid inclusions (NTDs distinguished by colour, all CTDs gray), with schematic of adjacent packing in capsid lattice. (**c**) Mature capsid structure after Pornillos et al. ^21^. NTDs of hexamer-associated CA shown in green, with NTDs of pentamer-associated CA shown in red (all CTDs gray), scale bar 20 nm.

HIV-1 capsid protein is mainly dimeric under physiological conditions, with CA monomers connected by a well-characterised CTD/CTD interface ^10^. Given suitable conditions in vitro, CA can spontaneously self-assemble into a wide variety of structures ^11-18^, but the prototypical mature virion contains a single conical capsid with a complex of viral RNA and nucleocapsid protein (NC) condensed within the broader terminus. In agreement with fullerene cone models ^14^, pentameric and hexameric CA oligomers have been identified as the basic components of mature-style CA lattice, with CA NTDs arranged into quasi-equivalent rings, ^15,19-21^ and with pentamers believed be present in regions of higher local curvature in the capsid lattice.^21,22^ Although the NTD and CTD structures of CA are well conserved, a flexible inter-domain linker region allows CA in solution to dynamically transition between conformations that are compatible and incompatible with mature lattice ^23^.

Controlled study of HIV-1 capsid assembly is complicated by the inherent variability of virions: non-trivial differences in the average radius (≈ 630 ± 50 Å) and Gag content (≈ 2400 ± 700 molecules) are reported in vivo ^24-26^, and these values bound a significant potential range of CA concentrations. The large numbers of Gag, in addition to viral RNA and sundry molecules sequestered from the host cell ^27,28^, ensure the virion interior is a highly crowded environment with internal organic mass densities as high as ≈ 300 mg/mL ^24^. Molecular crowding effects are known to influence protein behaviours ^29^ and yet, remarkably, virion maturation consistently produces similar capsid morphologies. The relatively fast maturation process ^30,31^ makes in vivo study of capsid self-assembly pathways difficult. Nonetheless, Woodward et al.^32^ show capsid formation proceeding via a hook-shaped precursor to the broad end of the capsid, in contrast with models of capsid assembly that begin with the narrow end of the cone ^33^.

As the average virion contains approximately twice the CA present in a mature capsid, a capsid exists in equilibrium with a relatively high concentration of solution-state CA. Viral infection requires the transfer of virion contents into a target cell, producing a rapid dilution of this CA solution. Rapid dilution destabilises CA lattice in vitro under physiological salt concentrations ^12^, but the potential significance of this effect on capsid stability is unclear.

Computer simulations of CA self-assembly offer a valuable complement to other experimental techniques. Atomic-resolution molecular dynamics (MD) simulations of pre-assembled capsids have been reported for HIV-1 ^22^, but the long time stability of such MD models has not been demonstrated and their great computational expense prevents such models being used to examine capsid self-assembly. Coarse-grained (CG) models using simpler molecular representations ^34,35^ provide an appealing alternative: for example, extensive CG simulations of icosahedral capsid self-assembly have been reported but the small, well-defined end products are in striking contrast to the structural pleomorphism observed with HIV-1 CA. Non-equilibrium CG simulations featuring irreversible subunit binding can generate capsid-relevant morphologies ^36,37^, but may struggle to represent any dynamic reorganization that might occur at the leading edges of lattice growth. The CG self-assembly of CA dimers was explored using Monte Carlo ^38^ and molecular dynamics simulations ^34^, but these studies allowed CA motion only on 2-D or quasi-2D surfaces.

In this work, we consider 3-D self-assembly behaviours of CG HIV-1 capsid protein under various conditions relevant to the viral lifecycle. In particular, we examine the effects of CA concentration and molecular crowding on the generation of mature CA lattice, the influence of dynamic transitions between populations of CA that are compatible and incompatible with mature lattice formation, and the effects of rapid CA dilution on previously self-assembled CA lattice structures. In total, we present the results of 51 separate CG simulations with general trends across the data sets examined.

## Results

### The effects of protein concentration and molecular crowding on the self-assembly ofCG capsid protein

As noted, there is a dynamic population of CA conformations dictated by the flexible linker between the NTD and CTD. For our simulations, the equilibrium NTD-CTD conformation was based on that observed in the curved hexagonal lattice within CA tubes (Figure S1). CGMD simulations were performed (see Methods and Supporting Information Fig. SI 1A-C) using CA concentrations [CA] of 1 mM, 2 mM, 3 mM, and 4 mM (with [CA] in a typical virion estimated as ≈ 3.8 mM) ^24^. For each [CA] studied, inert CG crowding agent densities *ρ*_*CR*_ from 0 mg/mL to 200 mg/mL were present with increments of 50 mg/mL to study the effects of molecular crowding on CG self-assembly ^12^. A single simulation was performed for each data point for 2 x 10^8^ MD time steps, with results summarised in Table 1 (full data in Supporting Information, Fig. SI 2).

**Table 1.** CG self-assembly for varying [CA] and inert crowding levels. Triangles (**Δ**) indicate simulations producing only steady-state CA trimer-of-dimers populations (average trimer count listed, standard deviation in parentheses). Entries marked ❖❖ denote production of multiple lattice regions. Hexamer (bold typeface) and pentamer (italics) counts shown after 2x10^8^ CGMD time steps.

| Inert crowding<br>level $\rho_{CR}$ / mg/mL | CA concentration / mM | | | |
| --- | --- | --- | --- | --- |
|  | 1 | 2 | 3 | 4 |
| 0 | $\Delta$ 0.36 (0.63) | $\Delta$ 2.98 (1.67) | $\Delta$ 10.47 (3.22) | $\Delta$ 25.49 (5.29) |
| 50 | $\Delta$ 0.57 (0.75) | $\Delta$ 5.41 (2.20) | $\Delta$ 18.74 (4.03) | $\Delta$ 42.11 (6.48) |
| 100 | $\Delta$ 1.32 (1.09) | $\Delta$ 10.73 (3.14) | $\Delta$ 33.19 (6.01) | $\diamond\diamond$ **68**, 7 |
| 150 | $\Delta$ 3.38 (1.72) | $\Delta$ 21.02 (4.23) | $\diamond\diamond$ **78**, 3 | $\diamond\diamond$ **140**, 6 |
| 200 | $\Delta$ 6.90 (2.57) | $\diamond\diamond$ **52**, 3 | $\diamond\diamond$ **96**, 16 | $\diamond\diamond$ **146**, 20 |

For [CA] = 1 mM, no stable self-assembly of mature-style capsid lattice occurred for any level of crowding studied. Instead, populations of metastable “trimer-of-dimers” structures ^34^ rapidly formed (Fig. 2 a,b), with average trimer populations increasing with *ρ*_*CR*_. A similar trend was observed for [CA] = 2 mM until 200 mg/mL of crowder was present, at which point the self-assembly behaviour changed dramatically. Rather than generating a steady-state population of trimers, as was apparently the case for *ρ*_*CR*_ = 150 mg/mL, a new self-assembly regime emerged: the steady production of mature-style CA lattice with quasi-equivalent pentamers and hexamers. Similar effects were observed for [CA] of 3 mM and 4 mM, with metastable trimers generated below the onset of CA lattice growth at 100 < *ρ*_*CR*_ ≤ 150 mg/mL (for [CA] = 3 mM) and 50 < *ρ*_*CR*_ ≤ 100 mg/mL ([CA] = 4 mM). The transition from producing only trimers to the nucleation and growth of mature-style lattice was thus a function of both [CA] and *ρ*_*CR*_, with higher [CA] requiring lower *ρ*_*CR*_ to induce lattice growth. Differences of ≤ 50 mg/mL in *ρ*_*CR*_ therefore led to markedly different behaviours for [CA] ≥ 2 mM.

**Figure 2.**
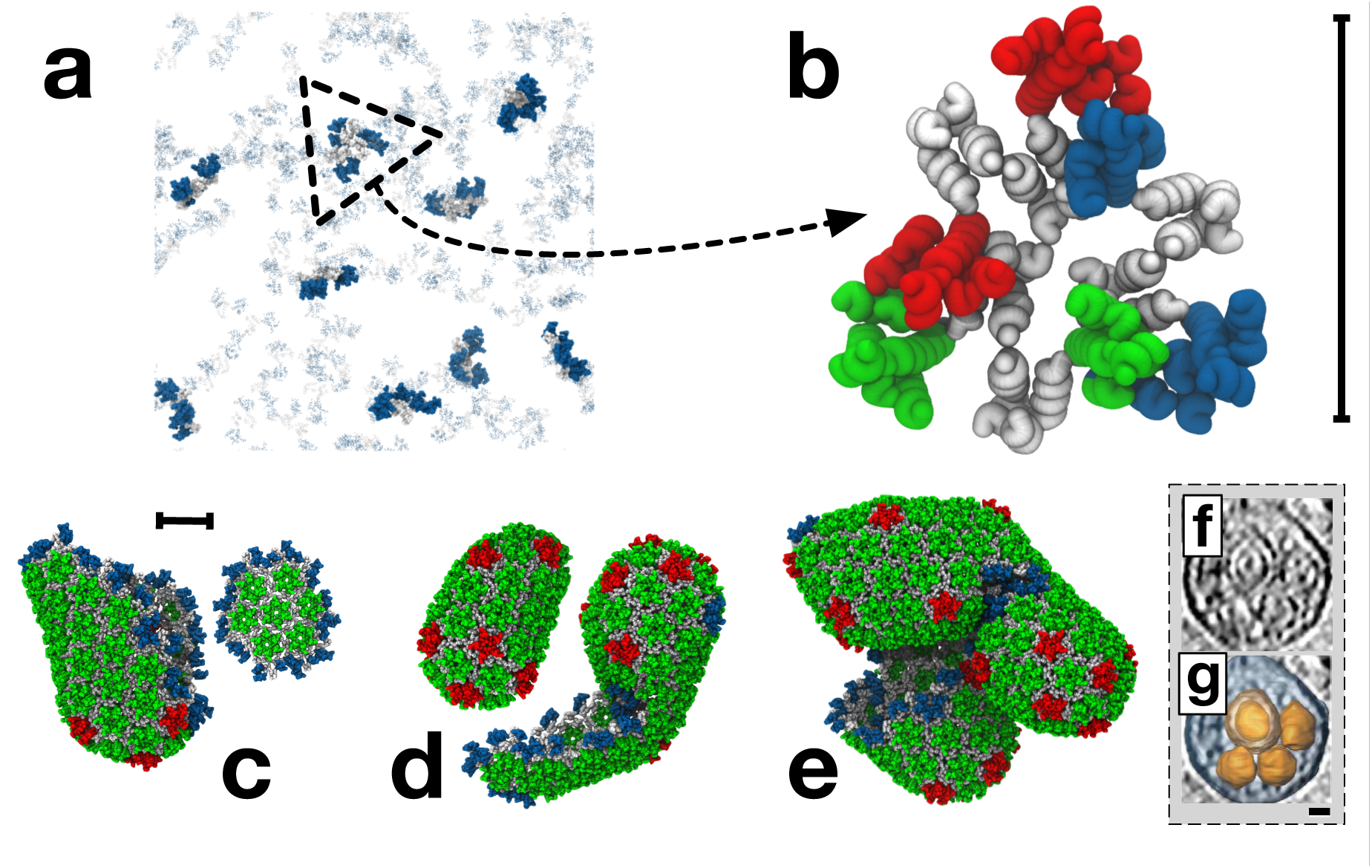
Dimers of the CA protein are the units of self-assembly of HIV-1 capsid-like CG structures, (**a**) Simulation snapshot of a population of metastable trimer-of-dimers. CA NTDs are blue, with CTDs gray (non-aggregated CA shown transparent for clarity), (**b**) Detail of a CG trimer-of-dimers, where “edges” are CA dimers, with two NTDs per triangle “vertex”. NTDs coloured by dimer for clarity, scale bar 10 nm. Final simulation snapshots for *ρ*_*CR*_ = 200 mg/mL and [CA] = 2 mM (**c**), 3 mM (**d**) and 4 mM (**e**) are presented with NTDs coloured by monomer presence in a trimer (blue), pentamer (red) or hexamer (green). All CTDs gray, scale bar 20 nm. (f) Multiple aggregated capsid structures as revealed by electron cryotomography,^39^ with capsid structures highlighted in orange (**g**). Scale bar 20 nm, panels **f** and **g** adapted from Reference 42.

Where simulations generated only steady-state trimer populations, the steady state was rapidly attained (< 5 x 10^6^ MD time steps). For simulations producing mature-style CA lattice, the appearance of stable pentameric inclusions lags the production of stable hexamers (see Fig. SI 2). In every simulation producing mature-style lattice, nucleation and growth of multiple lattice regions occurred. For example, with [CA] = 2 mM and *ρ*_*CR*_ =200 mg/mL, two independent lattice regions were generated: an incomplete cone fragment, and a small array of hexamers (Fig. 2c). For [CA] = 3 mM and *ρ*_*CR*_ =200 mg/mL, several independent lattice region appeared, resulting in a pill-shaped structure with a closed lattice surface and the fusion of an incomplete pill-shape with a cylinder fragment (Fig. 2d). For [CA] = 4 mM and *ρ*_*CR*_ =200 mg/mL, multiple lattice regions fused into a semi-amorphous assembly (Fig. 2e), reminiscent of multiple capsid structure aggregations observed in electron cryotomograms (Fig. 2 f-g,).^39^ The CG model is thus capable of producing a wide variety of curvatures, as required by the continuous curvatures observed in pleomorphic HIV-1 capsids.

Given the molecular mass of CA ≈ 25 kDa, [CA] of 2 mM, 3 mM, and 4 mM correspond to CA mass densities of ≈ 50 mg/mL, ≈ 75 mg/mL and ≈ 100 mg/mL. The onset of mature-style lattice growth therefore occurred for total mass densities of between 200 mg/mL and 250 mg/mL (for [CA] = 2 mM), 175 mg/mL and 225 mg/mL ([CA] = 3 mM), and 150 mg/mL and 200 mg/mL ([CA] = 4 mM). Total non-lipid mass density of HIV-1 virions was estimated as ≈ 200 to 300 mg/mL ^24^, and so the CG model self-assembly initiated at mass density levels comparable to the virion.

These results indicate a pronounced sensitivity to both CA concentration and local molecular crowding for CG self-assembly. Well-regulated nucleation and growth pathways are therefore critical for capsid production, given the significant natural variability for HIV-1 virions ^24-28^.

### The effects of capsid protein concentration on CG self-assembly under constant levels of molecular crowding

The level of inert molecular crowding can significantly alter CG self-assembly behaviours. However, CA is also a self-crowding agent as it occupies volume in the simulations. Direct comparison between self-assembly behaviours of different CA concentrations with the same quantity of inert crowder is therefore non-trivial.

To examine the effects of CA concentration on CG self-assembly under constant molecular crowding, the initial CA content of identical systems (4 mM CA, *ρ*_*CR*_ = 200 mg/mL) was partitioned into fixed “active” ([CA]_*+*_) and “inactive” ([CA]_*-*_) populations. After this initial partitioning, CA dimers therefore remain either “active” or “inactive” for the duration of the simulation. This partitioning is motivated by the experimental results of Deshmukh et al. ^23^ that suggest there is a relatively small population of “assembly-competent” CA dimers in CA solution. Inactive CA was identical to active CA in all aspects in our CG model except for the removal of attractive interactions, and thus both types of CA occlude equivalent volume. By fixing [CA]_*+*_ between 250 μM and 4 mM (with increments of 250 μM), CG self-assembly was studied as a function of CA concentration alone while maintaining fixed, virion-relevant crowding levels. The formation of trimer-of-dimers, hexamer, and pentamer structures were recorded as a function of simulation time over 2 x 108 CGMD timesteps (one simulation per CA concentration), with selected results summarised in Fig. 3 (full data in Table 2 and Fig. SI 3).

**Figure 3.**
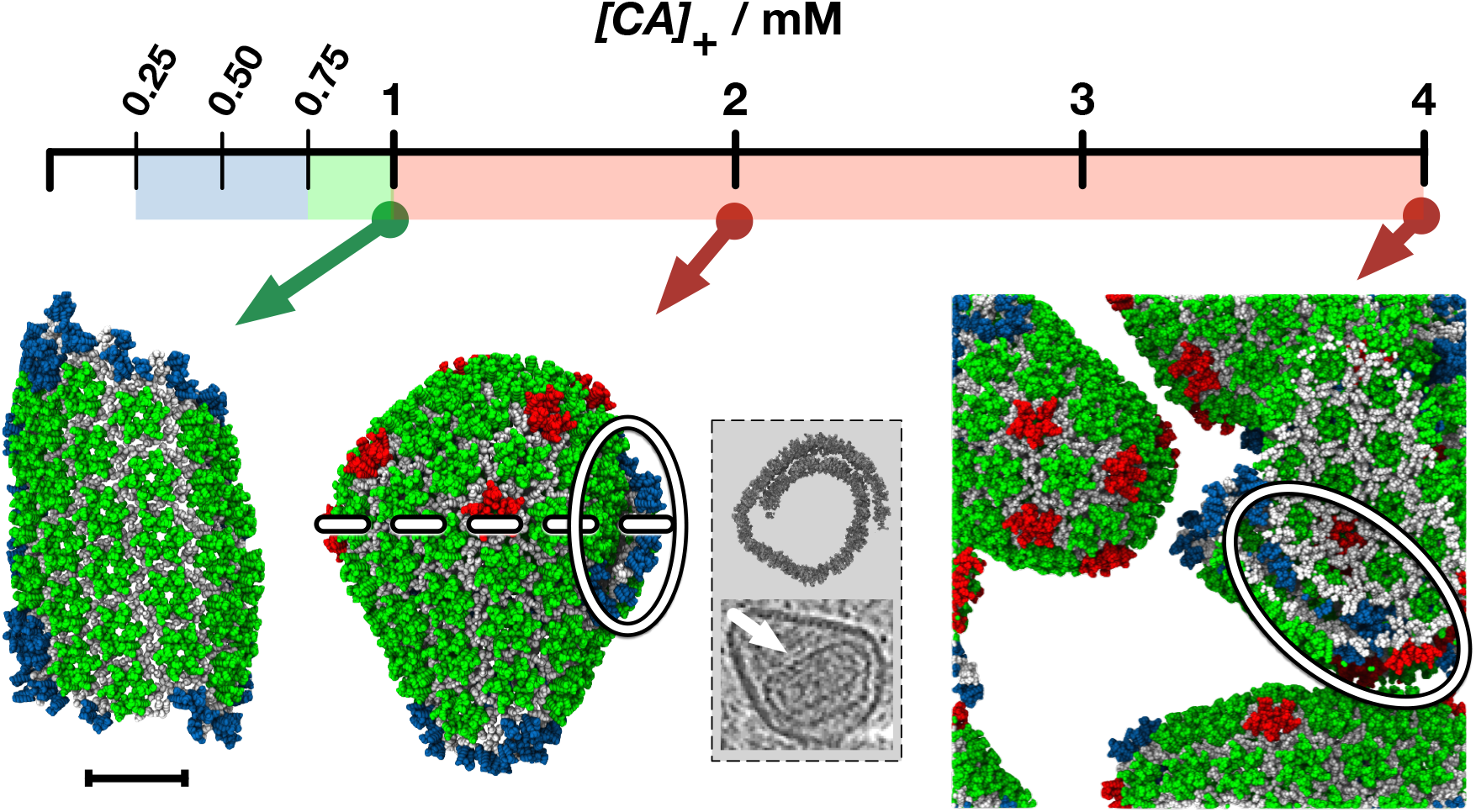
Only a narrow range (indicated in green) of CA concentrations results in the nucleation and growth of a single lattice region. CG self-assembly as a function of active CA concentration [CA]_*+*_ under fixed crowding conditions. Colour in the concentration bar indicates formation of only trimer-of-dimers structures (blue) and the nucleation and growth of single (green) or multiple (red) lattice regions. Example final structures are shown for [CA]_*+*_ = 1 mM, 2 mM and 4 mM (arrows). CA colour scheme as in Fig. 2, lamellar regions highlighted by ovals. Final structure for [CA]_*+*_ = 2 mM formed via two lattice regions fusing. Gray panel shows cross-sectional slices (not to scale) to illustrate lamellar lattice in structure for [CA]_*+*_ = 2 mM (top, cross sectional plane indicated by dashed white line) and an example lamellar CA lattice inside a virion from electron cryotomography (bottom, CA lattice indicated by white arrow). Final structures for [CA]_*+*_ = 4 mM wrap around periodic boundaries. Scale bar 20 nm.

**Table 2.**
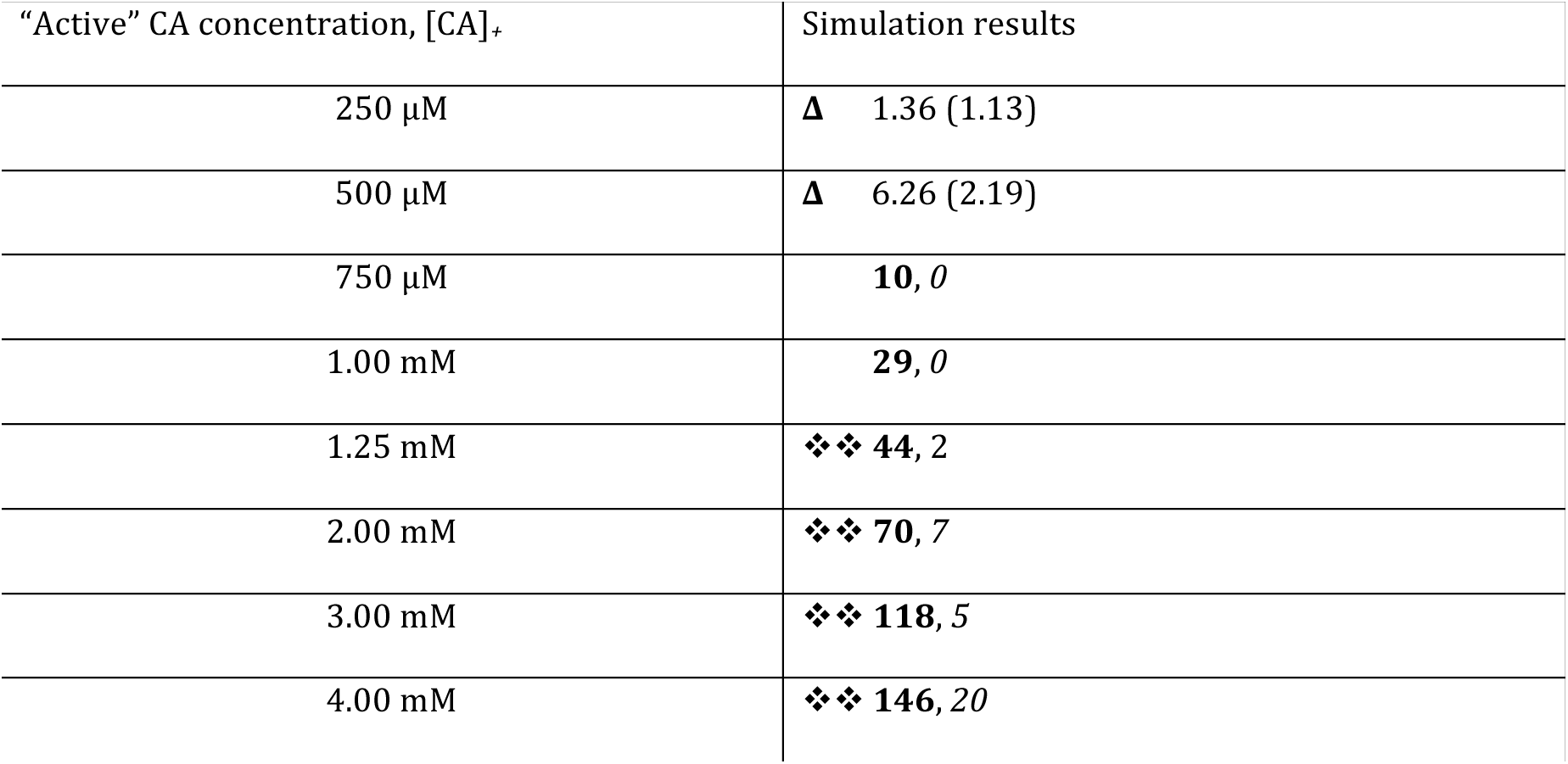
CG self-assembly for fixed “active” CA concentrations *[CA]*_*+*_ under constant crowding. Triangles (**Δ**) indicate simulations producing only steady-state populations of trimer-of-dimers structures (average trimer count listed, standard deviation in parentheses). Entries marked ❖❖ denote the production of multiple mature-style lattice regions. Final hexamer (bold typeface) and pentamer counts (italics) are shown after 2 x 10^8^ MD timesteps.

For 250 ≤ [CA]_*+*_≤ 500 μM, no generation of mature-style CA lattice occurred, but metastable trimer-of-dimer populations emerged. Elevating [CA]_*+*_to 750 μM produced an apparent initial steady-state trimer population before a single region of mature-style lattice nucleated and grew after ≈ 4 x 10^7^ MD timesteps. With [CA]_*+*_ = 1 mM, the nucleation and growth of a single lattice region began after ≈ 1 x 10^7^ MD timesteps. For both [CA]_*+*_ = 750 μM and [CA]_*+*_ = 1 mM, the lattice region adopted a cylindrical conformation as growth proceeded; although pentameric inclusions regularly formed at the growing edge of the lattice, the inclusions were transient and thus the lattice was essentially hexameric. For all [CA]_*+*_ ≥ 1.25 mM nucleation and growth of multiple lattice structures occurred, with regions of high local curvature around the pentamers. Also evident at higher [CA]_*+*_ were lamellar regions of CG lattice (Fig. 3) reminiscent of structures observed in electron cryotomograms. Lamellar and off-pathway capsid structures are indicated to be significantly more common in virion systems lacking RNA/NC complex,^40^ as is the case in these CG simulations.

The relatively controlled nucleation and growth for lower active CA concentration [CA]_*+*_ reveals specific details of CG lattice nucleation. Visual inspection of simulation trajectories indicate lattice nucleation occurring by the addition of CA dimers onto existing trimer-of-dimers structures, producing trimers with shared edges (Fig. 4) and leading to a central hexamer stabilised by a peripheral trimer “skirt” from which lattice growth proceeds. Metastable trimers therefore appear to be seeds for the nucleation and growth of CG lattice ^34^, rather than independent trimers aggregating. Transient structures along this pathway are [observed in simulations that do not produce mature lattice, but these intermediates are unstable and dissociate before nucleation of lattice growth.

**Figure 4.**
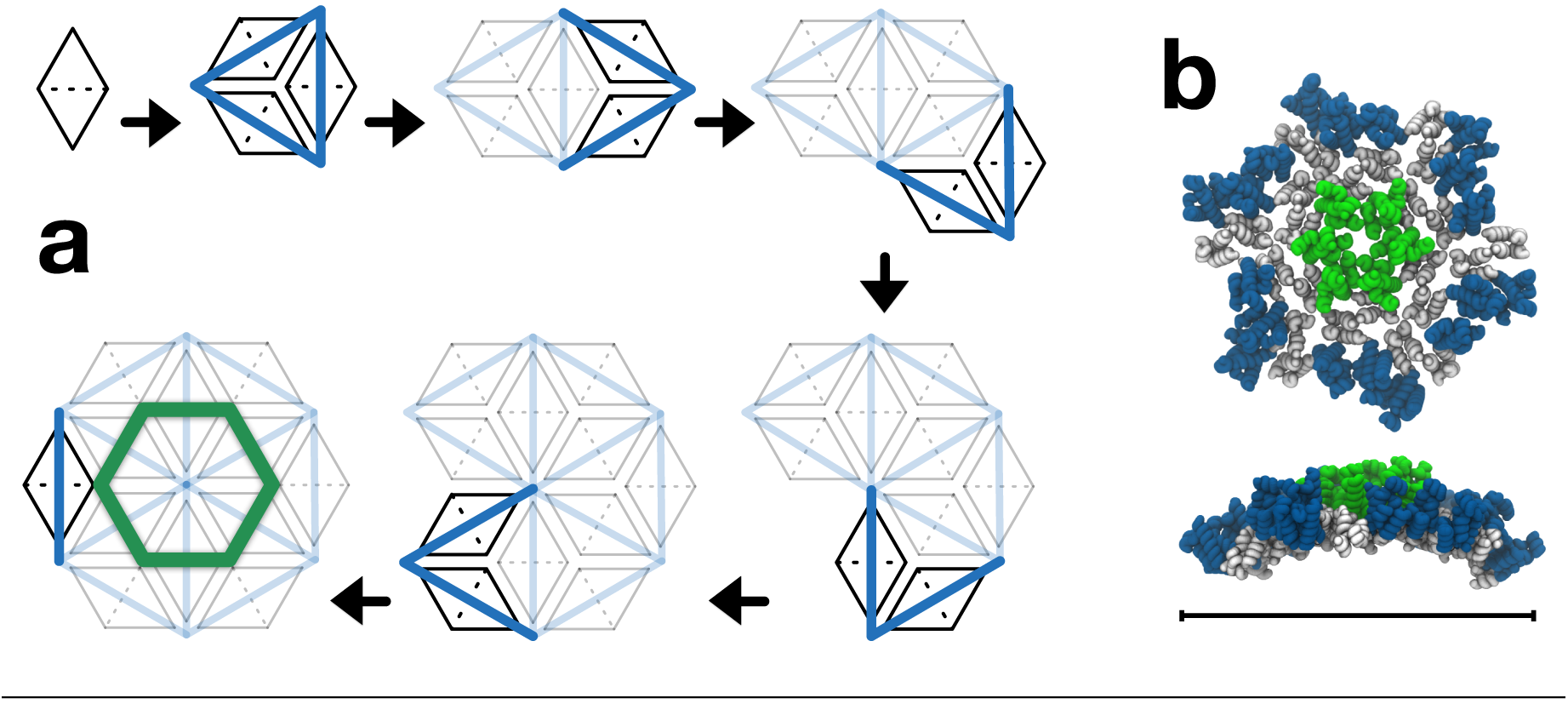
Putative assembly from 12 CA dimers that nucleates CG capsid lattice growth. (**a**) Reversible addition of CA dimers (CA monomers depicted as black triangles, CTD/CTD interface as a dashed line) onto existing aggregates produces trimers with shared edges, eventually generating a central hexamer stabilised by trimer “skirt”. (**b**) Front and side view of example structure from CG simulation, with mild innate curvature visible. Hexamer-associated NTDs are green, NTDs in trimer skirt blue, CTDs gray. Scale bar 20 nm.

Examination of the solution-state [CA]_*+*_ reveals that lattice growth continues below the level of solution-state [CA]_*+*_ required for initial nucleation (e.g. Fig. 5a), in agreement with standard nucleation and growth theories ^41^. As a region of mature lattice grows, there is also a reduction in the number of separate trimers in solution (e.g. Fig. 5b). Progressive reduction of not only the “available” capsid protein but also the number of trimers in solution (the potential nucleating seeds) can thus provide an elegant feedback mechanism to deter multiple capsid formation.

**Figure 5.**
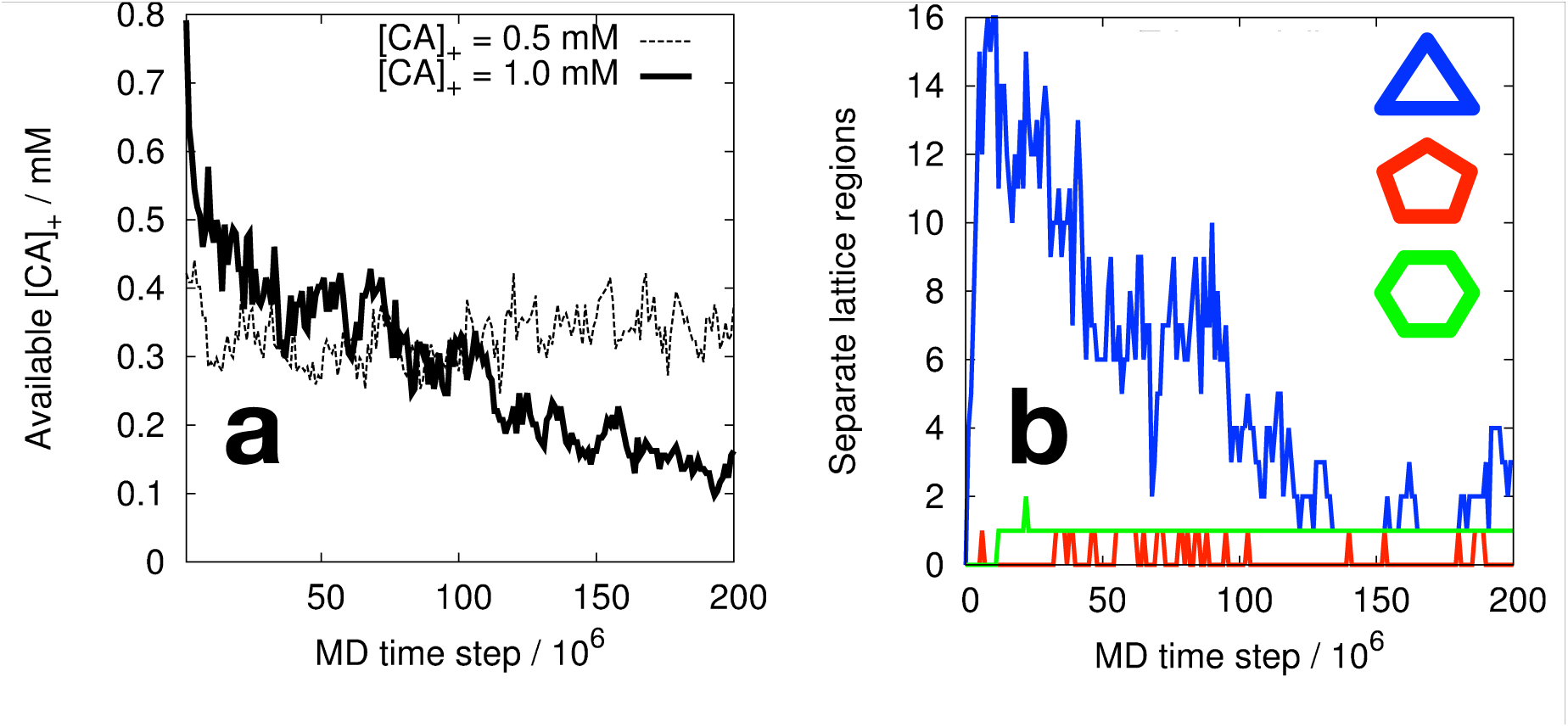
Example CG simulation data under fixed molecular crowding. (**a**) “Available” assembly-competent CA in solution for initial [CA]_*+*_ = 0.5 mM (resulting in no nucleation and growth of mature lattice) and initial [CA]_*+*_ = 1.0 mM (producing nucleation and growth of a single lattice region, see main text and Table 2). Lattice growth continues below the level of available [CA]_*+*_ required for nucleation on the same time scale. (**b**) Number of separate lattice regions containing key structural motifs for [CA]_*+*_ = 1 mM. The number of trimers in solution (blue curve) reduces significantly as a single region of lattice grows.

These results indicate a marked sensitivity of CG self-assembly processes to CA concentration under constant molecular crowding. Increases of as little as 250 μM in [CA]_*+*_ can drive the system from steady-state populations of metastable trimer structures ([CA]_*+*_ < 750 μM) into the nucleation and growth of single (750 μM ≤ [CA]_*+*_ ≤ 1 mM) or multiple lattice regions ([CA]_*+*_ ≥ 1.25 mM) over the time scales examined. Greater quantities of [CA]_*+*_ may encourage pentamer formation and/or stability over the course of the simulations: for [CA]_*+*_ ≤ 1 mM, no stable pentamers were detected (Table 2).

### The effects of CA NTD/CTD conformational freedom on CG self-assembly under constant protein concentration and molecular crowding

The NTD and CTD structures of CA are well conserved, but a flexible linker region allows significant conformational freedom in solution. Deshmukh et al. estimate that the solution-state CA with NTD/CTD arrangements compatible with mature lattice might be as low as ≈ 5% ^23^. Furthermore, this population is inherently dynamic, with correlation times for inter-domain motions estimated as between 2 ns and 5 ns ^23^. The CGMD simulations with fixed populations of “active” ([CA]_*+*_) and “inactive” ([CA]_*-*_) CA effectively describe a limiting case of this behaviour: [CA]_*+*_ corresponds to a population of CA whose instantaneous structure is compatible with mature lattice (and hence may directly self-assemble) and the fixed populations of [CA]_*+*_ and [CA]_*-*_ correspond to infinitely long correlation times for inter-domain motions (i.e., specific proteins remain in either assembly competent or incompetent conformations for the entire simulation).

The effects of interconversion between structural populations are now considered. Rather than initial fixing of [CA]_*+*_ and [CA]_*-*_ populations, all “free” CA in solution (i.e., CA which is not currently aggregated) is periodically identified and then randomly assigned to the [CA]_*+*_ or [CA]_*-*_ populations to maintain a specific target ratio of [CA]_*+*_ and [CA]_*-*_ in solution. The interval between random population assignments is thus a simplified proxy for time correlations in CA structural conformation: once assigned to either the [CA]_*+*_ or [CA]_*-*_ populations, a CA dimer remains “assembly competent” or “assembly incompetent” at least until the next random population assignment. Aggregated CA is not included in this process to avoid perturbation of existing aggregates, and to reflect CA conformational stability in mature CA lattice. Specific CA can disaggregate and, in the absence of re-aggregation, would be subject to the next random population switch. This approach allows significant control over the quantity of assembly competent CA in solution in order to mimic the coexistence of assembly competent and incompetent CA populations in solution.

For an identical baseline system (4 mM total CA, 200 mg/mL crowder), target “assembly competent” proportions [CA]_*%*_ of 2.5%, 5%, 10%, 25%, and 50% of solution-state CA were studied (one simulation per [CA]_*%*_ value). Random population switching was performed with intervals of 1 x 10^4^,1 x 10^5^, and 5 x 10^5^ CGMD time steps, corresponding to 0.1 ns, 1 ns, and 5 ns of CG simulation time respectively, but caution should be exercised in any direct comparison of CG timescales to those suggested by NMR experiments (see note regarding CG time scales in SI 4). Simulations were performed for 2 x 10^8^ CG MD time steps unless explicitly noted, with results summarised in Table 3 and Fig. 6 (full data in Fig. SI 4).

**Table 3.** CG self-assembly for varying [CA]_*%*_, (see main text) with 4mM total CAand 200 mg/mL of inert crowder. Triangles (Δ) denote steady-state trimer-of-dimer populations with no stable assembly of mature lattice (mean trimer count shown, standard deviation in parentheses). Entries marked ❖❖ indicate nucleation and growth of multiple lattice regions. Final hexamer (bold typeface) and pentamer count (italics) shown after 3 x 10^8^ MD timesteps unless otherwise noted in main text.

| Proportion of “active”<br>CA in solution, [CA]<sub>%</sub> | Switching interval |  |  |
| --- | --- | --- | --- |
|  | 1 x 10<sup>4</sup> time steps | 1 x 10<sup>5</sup> time steps | 5 x 10<sup>5</sup> time steps |
| 2.5% | $\Delta$ 0.13 (0.38) | $\Delta$ 0.33 (0.67) | $\Delta$ 0.66 (0.72) |
| 5% | $\Delta$ 0.20 (0.43) | $\Delta$ 0.83 (0.87) | $\Delta$ 1.63 (1.45) |
| 10% | $\Delta$ 0.39 (0.66) | $\Delta$ 2.43 (1.60) | **100**, *5* |
| 25% | $\Delta$ 1.99 (1.41) | $\diamond\diamond$ **82**, *7* | $\diamond\diamond$ **118**, *7* |
| 50% | $\diamond\diamond$ **90**, *4* | $\diamond\diamond$ **121**, *10* | $\diamond\diamond$ **128**, *13* |

**Figure 6.**
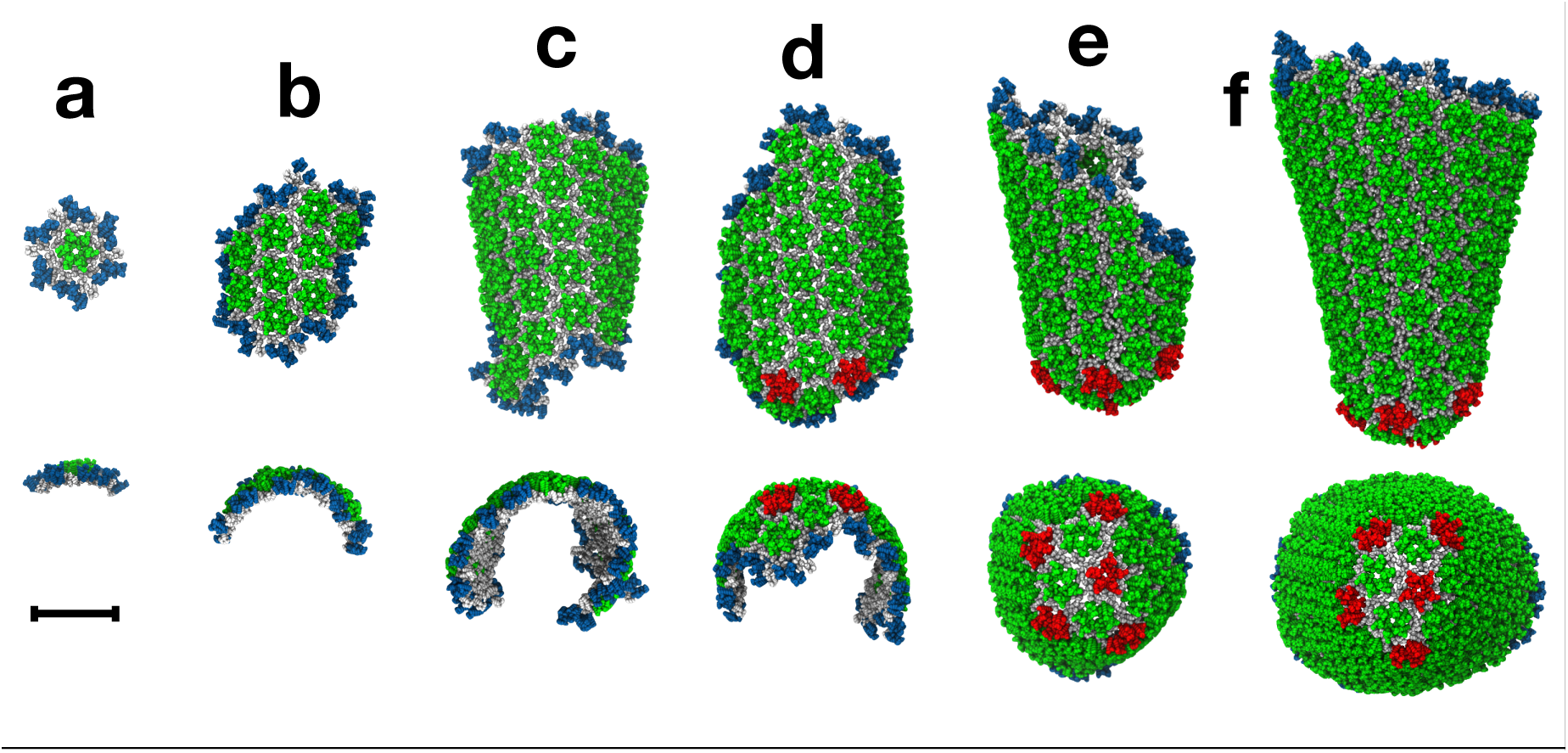
Steps in the assembly of the HIV-1 capsid by polymerization of CA dimers for [CA]_*%*_ = 10% and conformational switching interval of 5 x 10^5^ time steps (see main text). Simulation snapshots at 120 x 10^6^(**a**), 240 x 10^6^ (**b**), 440 x 10^6^ (**c**), 460 x 10^6^ (**d**), 600 x 10^6^ (**e**) and 1700 x 10^6^MD time steps (**f**) are shown with views perpendicular and parallel to the major structural axis. Colour scheme as in Fig. 2. Scale bar is 20 nm.

Apparent from Table 3 is the influence of conformational switching intervals on CG self-assembly. For the most rapid switching rate (interval = 1 x 10^4^ time steps), no self-assembly of mature-style CA lattice emerges until some 50% of the total CA in solution is considered to be in an assembly competent form: below this level, only metastable trimer-of-dimer structures appear. With a slower switching rate (interval = 1 x 10^5^ time steps), the transition from steady-state trimers into mature-style lattice occurs for 10% < [CA]_*%*_ ≤ 25%. For the slowest switching rate studied (5 x 10^5^ time steps), the transition occurs at 5% < [CA]_*%*_ ≤ 10%. For simulations producing only metastable trimers, faster switching rates reduces the average trimer population for identical [CA]_*%*_ (with an additional 1 x 10^8^ MD time steps performed to improve statistical estimates). These results suggest that correlation times for the NTD/CTD motions in capsid protein affect CG self-assembly, with faster switching between assembly competent and incompetent forms suppressing lattice formation for otherwise identical [CA]_*%*_.

When a target active proportion [CA]_*%*_ of 10% was combined with a switching interval of 5 x 10^5^ time steps, a cone-shaped structure reminiscent of a mature capsid was produced (Fig. 6). A small region of hexameric lattice initially formed, which curled into a semi-cylinder under growth. Stable pentamer incorporation appeared alongside higher local curvatures at one end of the structure, generating the narrow end of a cone. At this point lattice growth essentially stopped due to insufficient assembly-competent CA in solution to drive growth, as the system contained only 50% of the CA content of a typical virion.^24-26^ Although transient pentameric inclusions regularly formed at the exposed lattice edges, all permanent capsomer additions were hexameric after this point. It is interesting to note that pentamers were occasionally embedded slightly behind the growing edge of the lattice, but were replaced by hexamers in a process of local remodeling. The CG self-assembly can thus demonstrate a degree of “error correction”, provided the growing lattice edge does not advance too far beyond the pentamers. We note that although these conditions produced the assembly of a cone-shaped structure reminiscent of a mature capsid in this specific simulation, such results should not be considered to imply that these particular conditions are unique in producing this outcome.

### Stability and uncoating of self-assembled CG capsid lattice structures

Capsid assembly is critical for HIV-1 infectivity. However, capsid destruction is also crucial to the viral lifecycle: successful infection is sensitive to the specific time at which a capsid uncoats after cell entry, with both premature and delayed uncoating detrimental to viral replication ^6,42^. Assuming virion radius ≈ 630 Å ^24^, the transfer of virion contents into a cell of radius ≈ 10 μm ^43^ produces a rapid dilution of the CA solution around the capsid. To examine the effects of rapid dilution, simulations using previously self-assembled lattice structures were performed with [CA]_*%*_ = 0 to approximate the negligible effective concentration of CA in solution after virion contents are transferred into a cell. Importantly, this approach preserves the level of molecular crowding and conformational switching rate under which the lattice structures originally formed, allowing investigation of rapid dilution on several different structures that are otherwise stable under identical conditions. This avoids the use of a single model structure, whose natural stability may differ subtly under different test conditions even in the absence of rapid dilution‥These simulations therefore probe the effects of rapid dilution specifically, representing a cytoplasmic environment that is otherwise identical to the notional virion in which the structures were generated.

Maximally assembled systems (i.e., using [CA]_*%*_ = 50%) for the three conformational switching rates described previously were used as starting configurations for rapid dilution, with [CA]_*%*_ now set to 0% in each case. Fig. 7 presents the resultant time series of structural data from one rapid dilution simulation per starting conformation, with dissolution of CA lattice evident for all systems. Conformational switching intervals of 1 x 10^4^ time steps (Fig. 7a) and 1 x 10^5^ time steps (Fig. 7b) show a distinct and rapid destruction of any remaining lattice at ≈ 175 x 10^6^ and ≈ 550 x 10^6^ CGMD time steps respectively, indicating lattice dissolution is not a simple function of time (lattice structures at these points shown in Fig. 7, inset). Interestingly, the 5 x 10^5^ time step switching interval showed a long-lived plateau between ≈ 350 x 10^6^ and ≈ 800 x 10^6^ MD time steps corresponding to reorganization of the remaining lattice into a pill-like structure with closed surface (Fig. 7c). This enhanced stability is due to the absence of exposed lattice edges: CA in edge-free lattice requires significant reorganization of surrounding lattice components to escape, but can dissolve from exposed edges directly.

**Figure 7.**
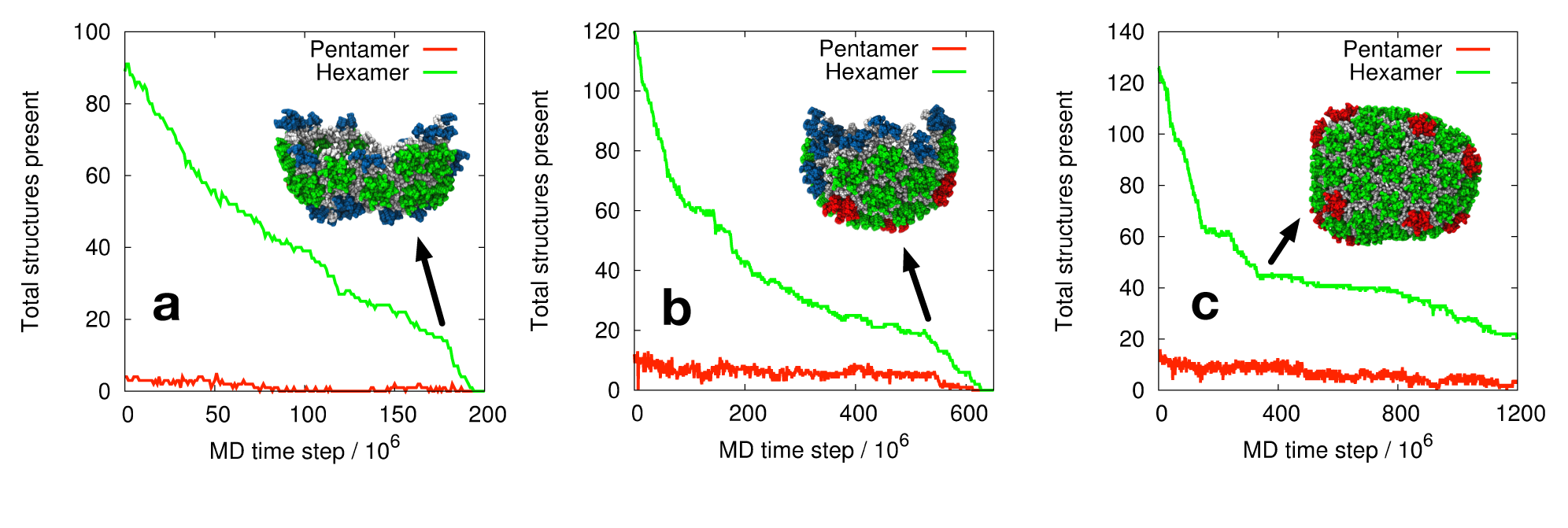
Disassembly of self-assembled CG CA lattice under simulated rapid dilution with constant molecular crowding. Conformational switching intervals of 1 x 10^4^ time steps (**a**), 1 x 10^5^ time steps (**b**), and 5 x 10^5^ time steps (**c**) are shown. Simulation snapshots (inset) correspond to the systems at≈ 175 x 10^6^ (**a**), ≈ 550 x 10^6^ (**b**) and ≈ 350 x 10^6^ (**c**) CG MD time steps. Colour scheme as in Fig. 2.

These results demonstrate the instability of CG capsid lattice under rapid dilution, even under conditions otherwise identical to those in which the lattice assembled. Furthermore, the results suggest that CA lattice structures with no exposed edges may be significantly more stable upon transfer into a cell.

## Discussion

Our study shows that CG self-assembly is sensitive to CA concentration, molecular crowding and NTD/CTD conformational correlation times. A single CG model generates surprisingly large morphological variety, encompassing the wide range of local curvatures required in the viral capsid ^15^. Evident in the results is a reversible multi-stage process triggering mature lattice self-assembly, ^44^ where metastable trimer-of-dimers structures rapidly emerge to act as potential seeds for the nucleation of mature lattice growth (Fig. 4). The steady-state population of trimers increases as a function of both concentration and molecular crowding, until suitable conditions encourage the generation of mature-style capsid lattice over the simulation lengths examined, progressively reducing both the solution-state CA and the population of independent trimers to deter the nucleation of additional lattice regions (e.g., Fig. 5). The transition between self-assembly regimes appears to be quite subtle, with relatively small changes to the molecular environment capable of producing different outcomes in agreement with the sensitivity displayed e.g. in previous studies of self-assembly for icosahedral capsid models. ^45-50^

The general effects of molecular crowding in our simulations are in agreement with those observed using other computational approaches to self-assembly. For example, high levels of crowding can reduce nucleation lag times to effectively zero (see e.g. Figs. SI 2-4), changing the basic nature of the self-assembly process from nucleation-limited to non-nucleation-limited. Schwartz and co-workers also noted such effects in discrete event simulations of icosahedral capsid self-assembly: ^48^ multiple parallel nucleation and growth events depleted the available capsid components, arresting complete capsid assembly and producing off-pathway growth. Similar kinetic traps and multiple nucleation was also observed in CG simulations of icosahedral capsid assembly.^45-50^

In the absence of RNA, CA is typically observed to assemble into tubes of variable diameter in vitro.^10,12,13,15-18^, albeit with occasional cone-shaped structures^12^ and other non-tubular morphologies also reported.^13,17^ These experiments are typically carried out with CA concentrations significantly below that expected in the virion, which is likely to assist in the generation of regular, purely hexameric cylinders via orderly nucleation and growth to help avoid off-pathway phenomena.

Sigmoidal “assembly curve” characteristics have been observed in experiment for capsid assembly systems in e.g. HIV-1^12^ and HPV-11. ^51^ Our results follow this general prediction, with lag periods for assembly that can be very short or effectively zero with CA concentration and/or significant crowding (see e.g. SI 2). We note that simulations lacking mature-style lattice production do not necessarily indicate that CG lattice growth cannot occur under these conditions, but rather that nucleation and growth was not detected over the simulation time scales examined. As the main focus of this study regards the nucleation and growth processes of capsid lattice, converged simulations (i.e., simulations that have reached some final steady state) do not offer significantly more information regarding these aspects. Nonetheless, examination of the number of structures relevant to mature lattice present in the simulations as a function of time (e.g., Figs. SI 2-4) indicates that many simulations are approaching the final plateau of a sigmoidal growth curve, and hence significant additional aggregation of capsid protein is unlikely.

Trimer-of-dimers structures have been reported previously in computer simulations ^34,38^, and have also been assumed as the fundamental building block in mathematical models of capsid cylinder growth ^52^. Our results suggest that, in isolation, these structures are metastable, and the addition of CA dimers onto these triangular templates (Fig. 4) provides the basic nucleation pathway for capsid assembly rather than the aggregation of independent trimer-of-dimer structures.

The regular appearance of transient pentamers at the unstable growing edge of CG lattice was observed, suggesting a natural aspect of lattice growth. If growth is relatively slow, local remodeling can remove the pentamers where the expanding lattice edge does not advance too far. For more rapid assembly (e.g., higher crowding or CA concentration) the lattice may advance too quickly for reliable remodeling, producing high local curvature via stably embedded pentamers. This process can explain why relatively low concentrations of CA typically produce cylindrical structures in vitro ^10,12,15-20^, and why increased CA concentration reduces cylinder length ^16^: early pentamer incorporation redirects the lattice growth, suppressing the formation of longer cylinders. The inherent instability of the lattice edge suggests that CG models that allow disassembly therefore more accurately reflect capsid assembly. ^36,37,44^

Switching between active and inactive CA populations in solution, akin to the conformational dynamism experimentally recorded in vitro ^23^, has pronounced effects on CG lattice self-assembly. Faster random mixing of the populations appears to suppress CG lattice production (Table 3), with a low instantaneous proportion of assembly-competent CA limiting uncontrolled nucleation and growth. This effect may be due to the properties of CG CA protein immediately after dissociation from a lattice region: with longer intervals between population switching, such CA may remain in an assembly-competent conformation long enough for protein/protein interactions to encourage direct re-association, but more investigation is required to fully elucidate this potential mechanism. In any case, dynamic conformational switching ensures that growth after nucleation is not prevented by the exhaustion of an otherwise limited quantity of suitable CA. This finding suggests that CG models incorporating this phenomenon may better represent a key aspect of HIV-1 capsid assembly.

It is interesting to consider HIV-1 virion maturation in light of these results. Although the prototypical mature virion contains a single capsid, multiple capsid structures are sometimes observed ^9,31-33,53^, both with and without RNA/NC encapsulation ^32^, suggesting a delicate balance in the virion. Multiple capsids appear correlated with larger virions ^33,53^, consistent with sensitivity of CG assembly to local conditions: lattice growth reduces capsid protein in solution, so larger virions (with larger numbers CA for the same overall CA concentration) could maintain CA levels above a critical nucleating value for longer periods after an initial lattice region forms. This increases the likelihood of additional nucleation events, compatible with observations that multiple cores are often comparable in size ^53^, which might suggest they nucleated in rather quick succession with no further nucleation occurring. The sensitivity of CG self-assembly to CA concentration and local molecular crowding, in combination with natural virion variability, may thus complicate previous analyses suggesting multiple core assembly is not a function of CA concentration ^53^.

Our CG simulations produced virion-relevant morphologies such as the extended cone-shaped structure shown in Fig. 6, including the initial generation of the narrow end of the cone. However, electron cryotomography suggests that capsid growth typically proceeds from a small region of CA lattice adsorbed to the RNA/NC complex, generating the broad end of the capsid first (although capsid growth also occurs without RNA/NC encapsulation). ^32^ The presence of such an RNA/NC complex in our simulations would prevent the high local curvature present in the narrow end of the capsid, and would thus select against the initial formation of this narrow end. Our simulations therefore do not fully recapitulate the true virion environment, but instead explore principles relevant to capsid self-assembly. Nonetheless, our results are in general agreement with models of capsid assembly that feature an initial small sheet of CA lattice that curls under growth, producing a cup shaped structure from which assembly proceeds.^32^

Our results suggest the importance of relatively slow, controlled growth of the HIV-1 capsid structure. Such a process allows significant relaxation of both the local and global structure during capsid growth, to allow the generation of a metastable capsid despite the apparently weak interactions between CA dimers.^11,12^ This phenomena suggests caution is appropriate when fitting atomic-resolution models into relatively low-resolution capsid data: for example, CTD/CTD dimer interfaces present in the atomistic MD simulations of HIV-1 viral capsids by Zhao and co-workers^43^ can display significant shearing and distortions from the expected structures^10,23^ (Fig. SI 5). It is possible that such deformations occur naturally in a viral capsid, but further experimental support is needed to clarify this situation.

The stability of CG lattice structures was sensitive to CA concentration in the supporting solution. Rapid dilution under otherwise identical conditions destabilised capsid lattice, indicating the metastability of CA structures in agreement with experimental observations ^12^. Similar effects have also been observed in simulations of icosahedral capsids,^54^ albeit these simulations did not examine this effect under constant crowding conditions. Sealed, edge-free CA lattices appear more resistant to this process (Fig. 7c). Cellular responses to HIV-1 include the weak binding of TRIM5α protein to capsids, forming a hexameric super-lattice on the capsid exterior ^55^. TRIM5α binding promotes rapid capsid uncoating ^42,56-61^, and while TRIM5α is digested by the proteasome ^62^ capsid protein does not appear to be destroyed in this manner ^56,57,61^. TRIM5α restriction is less efficient under proteasomal inhibition ^63-65^, and we speculate this somewhat counterintuitive data can be explained by TRIM5α inducing local faults in the capsid lattice: slight mismatches in CA and TRIM5α lattice spacing induces additional stress in the metastable CA lattice ^66^, with local tears providing an exposed edge from which CA can dissolve. Proteasomal degradation then ensures that a TRIM5α “scaffold” cannot exert any stabilizing influence on the overall capsid superstructure, enabling fragmentation of the capsid to further accelerate uncoating. Partial support for this hypothesis is offered by TRIM5α inducing only mild disruption of CA cylinders in vitro ^59^, but as these studies used above-physiological salt concentrations with no rapid dilution (conditions in which the CA lattice may be over-stable) ^12^ complete breakdown of CA lattice was not observed. Experimental data suggests HIV-1 capsids often feature exposed lattice edges, with an estimated 25% of cytoplasmic capsids featuring seams or holes large enough to allow the escape of GFP,^37^ and so TRIM5α could help to accelerate natural lattice dissipation under rapid dilution. For capsid structures lacking exposed lattice edges, however, TRIM5α would provide a powerful accelerator of uncoating. This process would offer a relatively simple description of the general effects of TRIM restriction on HIV-a capsids, but we note that the natural TRIM restriction processes could be significantly more complicated.

## Methods

### Model Details

Full CG model details are provided in SI 1. Briefly, CG capsid protein is modeled directly from experimental hexamer (PDB 3H4E) and pentamer (3P0A) data ^20,21^, and also heavily influenced by work from the same authors (B. K. G-P., M.Y.) with cylindrical CA assemblies in vitro ^14-16,19,20^. CA is represented as a dimer, the prevalent species in solution ^10,23^, which is required for CA lattice assembly ^11^. The NTD and CTD of a CA monomer are represented as independent stiff elastic network models (ENMs) comprising the C*α* atoms from CA alpha helices, which are well conserved across experimental structures ^34^. An additional weaker ENM connects the NTD and CTD in each monomer to provide limited inter-domain structural flexibility. The CTD/CTD dimer interface is represented by a stiff ENM based on PDB structure 2KOD ^10^ relevant for mature lattice ^23^. CG beads have excluded volume radii to deter unphysical overlaps as indicated by experimental structural data. Important CA protein/protein interfaces required for self-assembly ^9,10^ are represented by specific attractive interactions, with the locations of energy minima parameterized using experimental structures^20,21^ and attractive strengths chosen to allow CA self-assembly under the experimentally relevant mass densities and conditions we study here (SI 1). These attractive interactions can be dynamically enabled and disabled, allowing the generation of a fixed solution-state ratio of assembly competent and assembly incompetent populations as indicated by NMR data.^23^ Inert crowding agent is based on the excluded volume and relative mass of Ficoll 70 ^12^. The combination of internal NTD/CTD flexibility in CA monomers, “switching” capability, and freedom of motion in a 3-D simulation domain with explicit crowding molecules provides an advanced and relatively detailed molecular model with which to study CA self-assembly.

### Simulation details

All simulations performed with our UCG-MD code ^67^. Simulation domain was an 800 Å cube (≈ 50% of typical virion volume for computational efficiency, with max. 616 CA dimers present at [CA] = 4mM) with periodic boundaries. Temperature of 300 K maintained by Langevin dynamics with relaxation period 100 ps^-1^ to minimise influence of thermostat, integration timestep 10 fs. Visualizations of simulation data were created using VMD,^68^ with CG CA depicted as tubes connecting CG particles in each notional alpha helix to emphasize the local secondary structure of the CG model.

## Acknowledgements

This research was supported by National Institutes of Health grant P50-GM082545. The computations in this work are part of the Blue Waters sustained-petascale computing project, which is supported by the National Science Foundation (awards OCI-0725070 and ACI-1238993) and the state of Illinois. Blue Waters is a joint effort of the University of Illinois at Urbana-Champaign and its National Center for Supercomputing Applications. This work is also part of the “Ultra-Coarse-Grained Simulations of Biomolecular Processes at the Petascale” Petascale Computing Resource Allocation (PRAC) support by the National Science Foundation (award number OCI-1440027).

